# Dice-XMBD: Deep learning-based cell segmentation for imaging mass cytometry

**DOI:** 10.1101/2021.06.05.447183

**Authors:** Xu Xiao, Ying Qiao, Yudi Jiao, Na Fu, Wenxian Yang, Liansheng Wang, Rongshan Yu, Jiahuai Han

**Author notes:** Corresponding authors {, }. These authors have contributed equally to this work (Xu Xiao, Ying Qiao).

## Abstract

Highly multiplexed imaging technology is a powerful tool to facilitate understanding cells composition and interaction in tumor microenvironment at subcellular resolution, which is crucial for both basic research and clinical applications. Imaging mass cytometry (IMC), a multiplex imaging method recently introduced, can measure up to 40 markers simultaneously in one tissue section by using a high-resolution laser with a mass cytometer. However, due to its high resolution and large number of channels, how to process and interpret the image data from IMC remains a key challenge for its further applications. Accurate and reliable single cell segmentation is the first and a critical step to process IMC image data. Unfortunately, existing segmentation pipelines either produce inaccurate cell segmentation results, or require manual annotation which is very time-consuming. Here, we developed Dice-XMBD, a Deep learnIng-based Cell sEgmentation algorithm for tissue multiplexed imaging data. In comparison with other state-of-the-art cell segmentation methods currently used in IMC, Dice-XMBD generates more accurate single cell masks efficiently on IMC images produced with different nuclear, membrane and cytoplasm markers. All codes and datasets are available at https://github.com/xmuyulab/Dice-XMBD.

## 1. Introduction

Analysis of the heterogeneity of cells is critical to discover the complexity and factuality of life system. Recently, single-cell sequencing technologies have been increasingly used in the research of developmental physiology and disease [1, 2, 3, 4], but the spatial context of individual cells in the tissue is lost due to tissue dissociation in these technologies. On the other hand, traditional immunohistochemistry (IHC) and immunofluorescence (IF) preserve spatial context but the number of biomarkers is limited. The development of multiplex IHC/IF (mIHC/mIF) technologies has enabled the detection of multiple biomarkers simultaneously and preserve spatial information, such as cyclic IHC/IF and metal-based multiplex imaging technologies [5, 6, 7, 8]. Imaging mass cytometry (IMC) [6, 9], one of metal-based mIHC technologies, uses a high-resolution laser with a mass cytometer and makes the measurement of 100 markers possible.

IMC has been utilized in studies of cancer and autoimmune disorders [6, 10, 11, 12, 13]. Due to its high resolution and large number of concurrent marker channels available, IMC has been proven to be highly effective in identifying the complex cell phenotypes and interactions coupled with spatial locations. Thus, it has become a powerful tool to study tumor microenvironment and discover the underlying disease-relevant mechanisms [14, 15, 16, 17, 18, 19,20, 21]. Apart from using IMC techniques alone, several other technologies, such as RNA detection in situ and 3D imaging, have been combined with IMC to expand its applicability and utility [22, 23, 24, 25].

The IMC data analysis pipeline typically starts with single-cell segmentation followed by tissue/cell type identification [26, 27, 28]. As the first step of an IMC data processing pipeline, the accuracy of single-cell segmentation plays a significant role in determining the quality and the reliability of the biological results from an IMC study. Existing IMC cell segmentation methods include both unsupervised and supervised algorithms. Unsupervised cell segmentation, such as the watershed algorithm implemented in CellProfiler [26], does not require user inputs for model training. However, the segmentation results are not precise when cells are packed closely or they are in complicated shapes. To achieve better segmentation results, it is possible to use supervised methods with a set of annotated images with pixel-level cell masks to train a segmentation classifier. However, the manual annotation task is very time-consuming and expensive as well since it is normally done by pathologists or experienced staff with necessary knowledge in cell annotation. Particularly, for multiplexing cellular imaging methods such as IMC, their channel configurations including the total number of markers and their selections are typically study-dependent. Therefore, manual annotation may need to be performed repeatedly for each study to adapt the segmentation model to different channel configurations, which can be impractical.

To overcome this limitation, a hybrid workflow combining unsupervised and supervised learning methods for cell segmentation was proposed [18]. This hybrid workflow uses Ilastik [27], an interactive image processing tool, to generate a probability map based on multiple rounds of user inputs and adjustments. In each round, a user only needs to perform a limited number of annotations on regions where the probability map generated based on previous annotations is not satisfactory. CellProfiler is then used to perform the single cell segmentation based on the probability map once the result from Ilastik is acceptable. This hybrid workflow significantly reduces manual annotation workload and has gained popularity in many recent IMC studies [10, 13, 14, 17, 21]. However, the annotation process still needs to be performed by experienced staff repeatedly for each IMC study, which is very inconvenient. In addition, the reproducibility of the experimental results obtained from this approach can be an issue due to the per-study, interactive training process used in creating the single cell masks. Hence, a more efficient, fully automated single-cell segmentation method for IMC data without compromising the segmentation accuracy is necessary for IMC to gain broader applications in biomedical studies.

Convolutional neural networks (CNNs) have been successfully used for natural image segmentation and recently applied in biomedical image applications [29, 30, 31, 32]. CNN-based U-Net was developed for pixel-wise cell segmentation of mammalian cells [33]. It has been demonstrated that the U-Net architecture and its variants such as Unet++[34], 3D Unet [35] and V-Net [36] can obtain high segmentation accuracy. Motivated by the outstanding performance of using U-Nets for cell segmentation [37, 38, 39], we developed Dice-XMBD, a deep neural network (DNN)-based cell segmentation method for multichannel IMC images. Dice-XMBD is marker agnostic and can perform cell segmentation for IMC images of different channel configurations without modification. To achieve this goal, Dice-XMBD first merges multiple-channel IMC images into two channels consisting of a nuclear channel containing proteins originated from cell nucleus, and a cell channel containing proteins originated from cytoplasm and cell membrane. Channels of proteins with ambiguous locations are ignored by Dice-XMBD for segmentation as they contribute little to the segmentation results. Furthermore, to mitigate the annotation workload, we adopted the knowledge distillation learning framework [40] in training Dice-XMBD, where the training labels were generated using Ilastik with interactive manual annotation as a teacher model. We used four IMC datasets of different channel configurations to evaluate the performance of Dice-XMBD and the results show that it can generate highly accurate cell segmentation results that are comparable to those from manual annotation for IMC images from both the same and different datasets to the training dataset, validating its applicability for generic IMC image segmentation tasks.

## 2. Materials and methods

### 2.1 Overview of the pipeline

In Dice-XMBD, we used a U-Net based pixel classification model to classify individual pixels of an IMC image to their cellular origins, namely, nuclei, cytoplasm/membrane, or background. The pixel probability map produced by the classifier was then used by CellProfiler (version 3.1.0) to produce the final cell segmentation results (Figure 1). The pixel classification model was trained on IMC images with pixel-level annotations. To mitigate the annotation workload, Ilastik was used as the teacher model to produce the classification labels for training. Furthermore, to obtain a generic pixel classifier that can be used across IMC datasets of different channel configuration, channels of different proteins were combined based on their cellular origins into two channels, namely, nuclear and cell (membrane/cytoplasmic) channels, respectively. Channels of proteins without specific cellular locations were ignored by Dice-XMBD. The pixel classification model was then trained on the combined two-channel images. Likewise, the same preprocessing was used at the prediction stage to produce the two-channel (nuclear/cell) images for pixel classification. Of note, although the prediction may be performed on images with different markers, the channels were always combined based on their origins so that pixel classification was performed based on the two channels of putative protein locations rather than channels of individual proteins.

### 2.2 Training and evaluation datasets

We used four IMC image datasets in this study. BRCA1 and BRCA2 [18] contain 548 and 746 images from patients with breast cancer with 37 and 35 markers respectively. T1D1 [10] and T1D2 [11] contain 839 and 754 images from patients with Type I Diabetes with 35 and 33 markers, respectively. Dice-XMBD was trained on a subset of BRCA1 dataset (*n* = 348) with 200 held-out images reserved for validation and test. To test the generalization ability of Dice-XMBD, we also tested the trained model on the other three independent IMC datasets (BRCA2, T1D1 and T1D2).

### 2.3 Generating groundtruth cell masks

The groundtruth cell masks for training were generated using Ilastik and CellProfiler. We used the smallest brush size (1 pixel) in annotating the image to avoid annotating a group of neighboring pixels of different classes. The annotation was performed in an interactive manner, where the random forest prediction model of Ilastik was updated regularly during annotation to produce an uncertainty map indicating the confidence level of the classification results produced by the prediction model. The annotation was then guided by the uncertainty map to focus on the regions with high uncertainty iteratively until the overall uncertainty values were low except for regions of which the boundaries were visually indistinguishable.

The initial annotation was performed on a randomly selected subset of the dataset. After the initial annotation, we loaded all the images from the dataset into Ilastik to calculate their uncertainty maps, and then selected those with the highest average uncertainty values for further annotation. This process was iterated until the uncertainty values of all images converged, i.e., did not significantly decrease for three iterations.

In the end, we annotated 49 images in BRCA1 to train the model in Ilastik. We then imported all the images of the BRCA1 dataset into Ilastik for batch processing and export their corresponding pixel classification probability maps for training Dice-XMBD. The probability maps were further input to CellProfiler to produce the “groundtruth” cell segmentation. In CellProfiler, we used the “IdentifyPrimaryObjects” module to segment the cell nuclei and used the “IdentifySecondaryObjects” to segment the cell membranes using the propagation method. The output masks from CellProfiler are regarded as “groundtruth” cell segmentation of the dataset for performance evaluation.

We also generated the groundtruth cell masks of the other three datasets by the same iterative procedure separately for testing the generalization ability of Dice-XMBD. During the process, 72 images in BRCA2, 39 images in T1D1, and 67 images in T1D2 were manually annotated.

### 2.4 Training the U-Net cell segmentation model

#### 2.4.1 Image preprocessing

The input IMC images were firstly preprocessed by hot pixel removal, dynamic range conversion, normalization, and image cropping/padding into fix-sized patches. First, we applied a 5 *×* 5 low-pass filter on the image to remove hot pixels. If the difference between an image pixel value and the corresponding filtered value was larger than a preset threshold (50 in our experiments), the pixel would be regarded as a hot pixel and its value would be replaced by the filtered value. As the dynamic range of pixels values differs among IMC images of different batches and different channels, we further min-max normalized all images to [0,255] to remove such batch effect as follows:

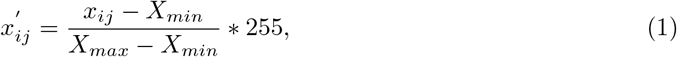

where *x*_*ij*_ is the pixel value in one channel, *X*_*max*_ and *X*_*min*_ denote the maximum and minimum values in the channel. Of note, as the pixel values in IMC images have a high dynamic range, transforming the pixel values from its dynamic range to [0, 255] would suffer from detail suppression by one or few extremely large values. Therefore, we thresholded the image pixel values at 99.7% percentile for each image before normalization.

Finally, we merged all the nuclear channels into the nuclear channel, and membrane/cytoplasmic channels into the cell channel, by averaging on all channel images with pre-selected sets of protein markers, respectively. We converted the merged two-channel images into patches of 512 *×* 512 pixels. For images or boundary patches that are small than target patch size, we set the pixel values of both channels to 0 and set the pixel type as background for padding.

#### 2.4.2 Data augmentation

Data augmentation is an efficient strategy to reduce overfitting and enhance the robustness of the trained models especially when training data is insufficient. We applied the following data augmentation methods on input images.

First, photometric transformations including contrast stretching and intensity adjustments were used. For contrast stretching, we changed the level of contrast by multiplication with a random factor in the range of [0.5, 1.5]. Similarly, for intensity adjustments we changed the level of intensity by multiplication with a random factor in the range of [0.5, 1.5]. Geometric transformations including image flipping and rotation were used. For flipping we implemented random horizontal or vertical flipping. For rotation, the rotating angle is randomly distributed in the range of [-180, 180]. Note that geometric transformations were applied to pairs of input and output images of the network. We also injected random Gaussian noise to the two input channels of the input images. Examples of data augmentation are shown in Supplementary Figures 1 and 2.

**Figure 1:**
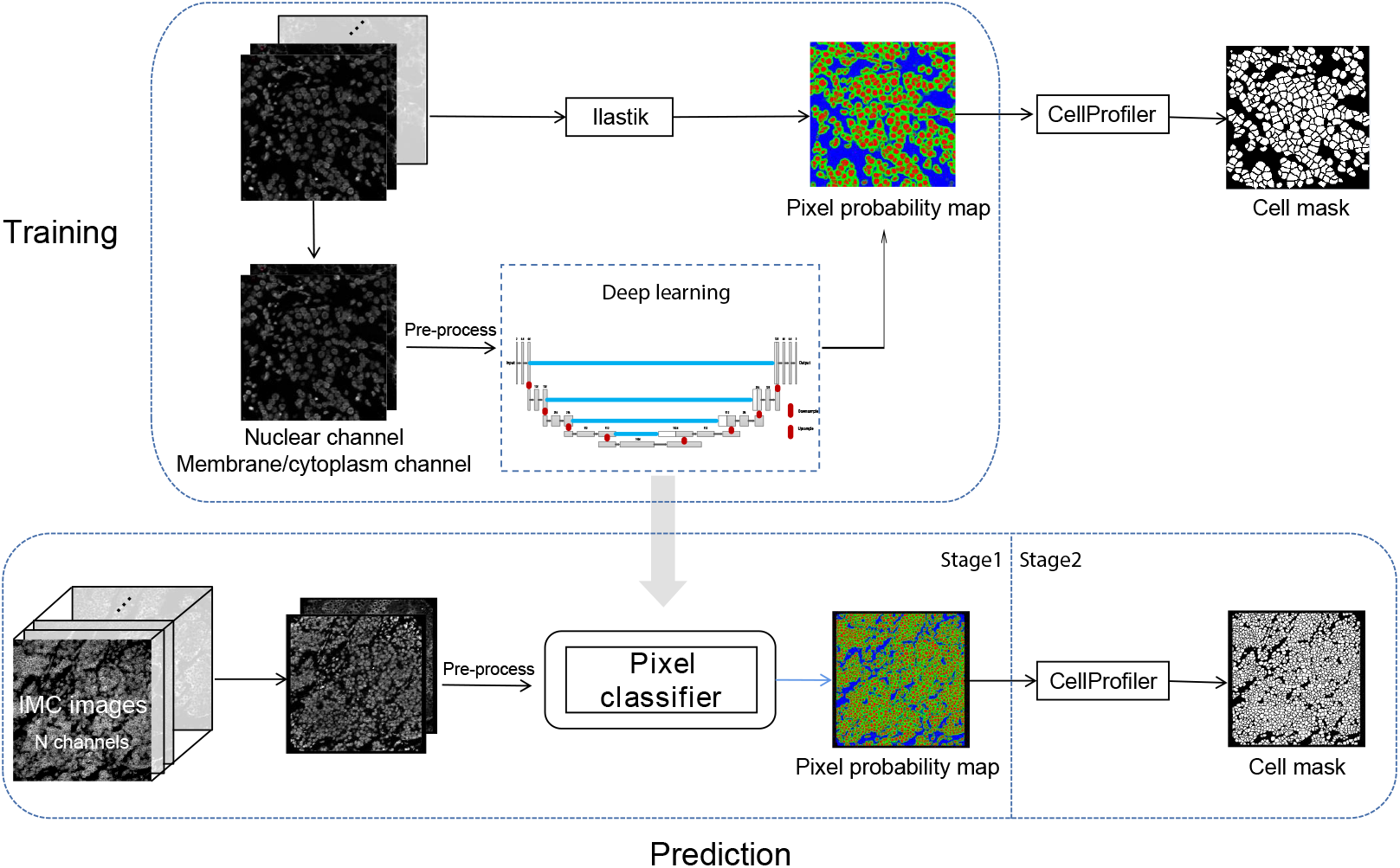
Dice-XMBD workflow. IMC images are combined into 2-channel images containing membrane/cytoplasm and nuclear proteins expression information. In stage 1, the pixel probability maps of given 2-channel images are predicted using a semi-supervised learning model based on Unet architecture. The training data were generated from Ilastik by using human annotations. In stage 2, the cell segmentation masks are generated from the pixel probability maps using the propagation method in CellProfiler.

**Figure 2:**
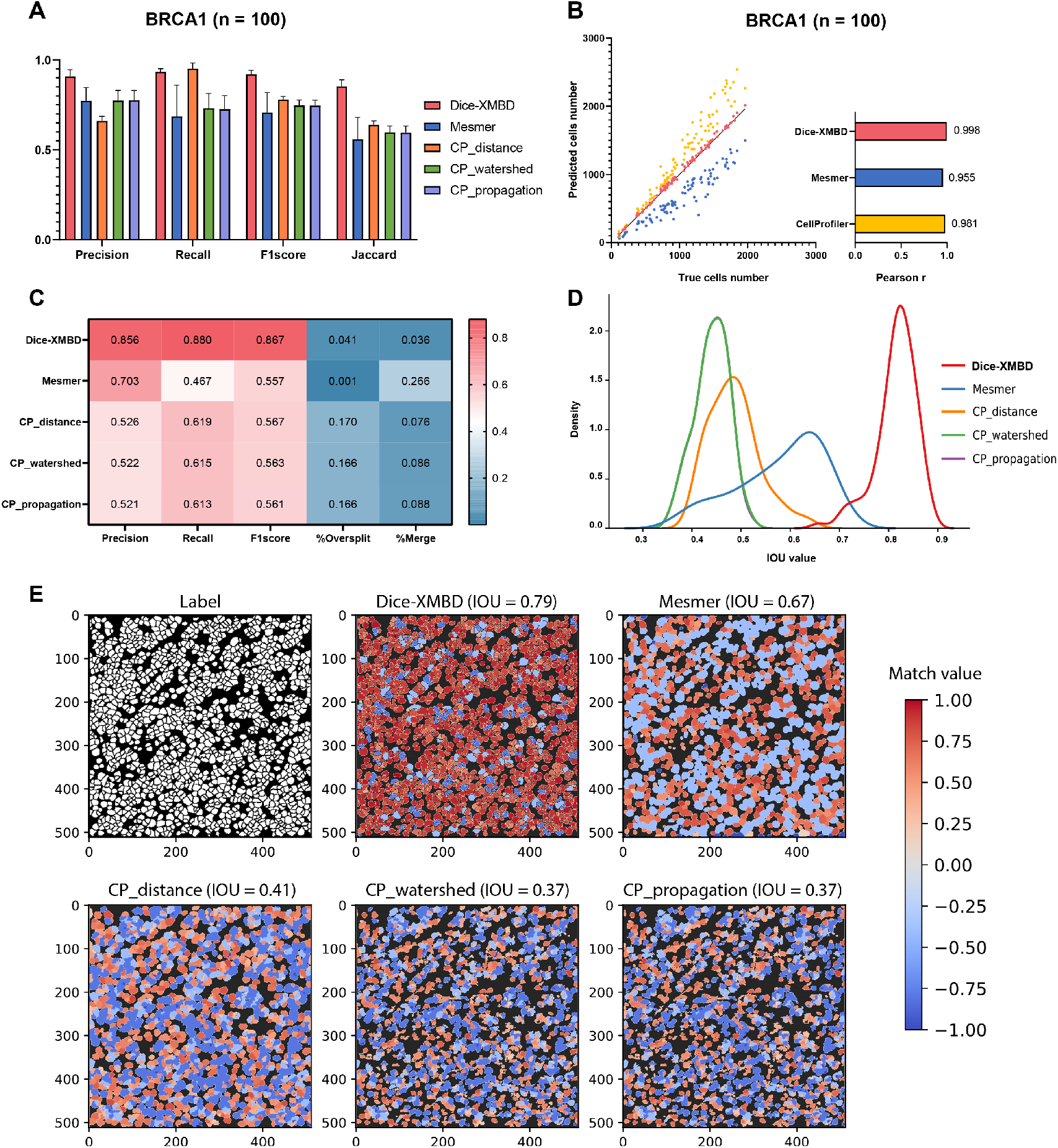
Dice-XMBD enables automatic single cell segmentation on dataset BRCA1. **(A)** Pixel prediction performance comparisons of Dice-XMBD, Mesmer and CellProfiler (CP-distance, CP-watershed, CP-propagation). All data in bar plots are presented as mean values +/-SD. **(B)** Pearson correlations between the number of predicted cells and labelled cells per image. The number of cells predicted from three cell segmentation methods implemented in CellProfiler are the same (represented as CellProfiler). **(C)** Cell prediction performance of five benchmarked methods. The percent of oversplits and merge errors in predictions are denoted as %Oversplit and %Merge. **(D)** Density plot shows the distribution of mean IOU values of matched cells per image. **(E)** An example of labeled and predicted single cell mask from benchmarked methods. Match value represents IOU value for one-to-one cell pair found in label and prediction, -0.4 and -0.8 for merged cells (multiple true cells are assigned to one predicted cell) and split cells (multiple predicted cells are matched to a true cell), and -1 for cells are not in any of above situations. Numbers in brackets of each method indicate mean of IOU values of all matched cell pairs.

#### 2.4.3 Constructing a pixel classification model

The U-Net pixel classification network is an end-to-end fully convolutional network and contains two paths. The contracting path (or the encoder) uses a typical CNN architecture. Each block in the contracting path consists of two successive 3 *×* 3 convolution layers followed by a Rectified Linear Unit (ReLU) activation and a 2 *×* 2 max-pooling layer. This block is repeated four times. In the symmetric expansive path (or the decoder), at each stage the feature map is upsampled using 2 *×* 2 up-convolution. To enable precise localization, the feature map from the corresponding layer in the contracting path is cropped and concatenated onto the upsampled feature map, followed by two successive 3 *×* 3 convolutions and ReLU activation. At the final stage, an additional 1 *×* 1 convolution is applied to reduce the feature map to the required number of channels. Three channels are used in our case for cell nuclei, membrane, and background, respectively. As we output the probability map, the output values are converted into the range of [0, 1] using the Sigmoid function.

#### 2.4.4 Loss function

We take the binary cross-entropy (BCE) as the loss function which is defined as:

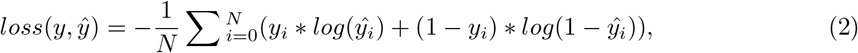

where *N* represents the total number of pixels in an image, *y*_*i*_ denotes the ground truth pixel probability and *ŷ*_*i*_ denotes the predicted pixel probability. The cross-entropy loss compares the predicted probabilities with the ground truth values. The loss is minimized during the training process.

### 2.5 Model evaluation

To evaluate pixel-level accuracy, we calculated the true positive and false positive based on the binary cell masks. In a binary cell mask, 1 represents cell boundary and 0 denotes cell interior or exterior. For every pixel in an image, true positive (TP) and true negative (TN) mean that the predicted pixel classification is the same with its label, while false positive (FP) and false negative (FN) mean that a pixel is misclassified as cell boundary or cell interior/exterior, respectively. To evaluate model performance at cell-level, we first calculated the intersection over union (IOU) on cells from predicted and labeled cell masks to determine if they are the same cell from different cell segmentation. After filtering cell matches with IOU below 0.1, if a predicted cell only finds one true cell, the cell is segmented accurately (same as TP). If a true cell cannot find a predicted cell, the cell is denoted as FN. Also, there exist some predicted cells which are assigned to the same true cell, we consider this situation as a spit error. If multiple true cells are matched to a same predicted cell, we consider those predicted cells as merge errors. Four standard indices are measured as below.

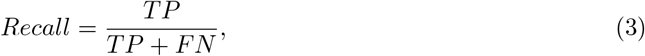

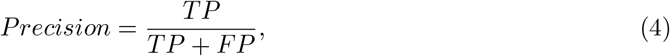

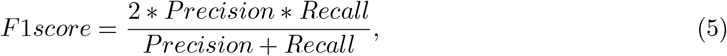

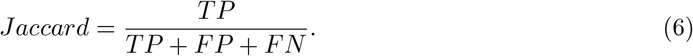

## 3. RESULTS

### 3.1 Dice-XMBD enables automatic cell segmentation

We trained a deep learning model with the above-described U-Net architecture using the BRCA1 dataset with 348 images as training set and 100 images as validation set. A complete held out test set with 100 images was used to test model performance within one dataset. We further applied the trained model directly on the other three IMC image datasets to evaluate the cross-dataset performance of the model. To evaluate the model performance, we computed standard indices (Recall, Precision, F1-score, and Jaccard index) for both pixel-level and cell-level accuracies (see Methods for more details).

We compared Dice-XMBD with a generic whole-cell segmentation method across six imaging platforms, Mesmer [41], which used a deep learning-based algorithm trained on a large, annotated image dataset to segment single cells and nuclei separately. A trained Mesmer model was tested with combined nuclear and cell channels which is the same as the input to Dice-XMBD. Meanwhile, we used three commonly used segmentation methods implemented in CellProfiler with default parameters: distance, watershed, and propagation. These methods first find nuclei as primary objects, and then the membrane proteins are added together into an image as input to recognize cells. The distance method does not use any membrane proteins information and simply defines cell membrane by expanding several pixels around nuclei. The watershed method computes intensity gradients on the Sobel transformed image to find boundary between cells [42], while the propagation method defines cell boundary by combining the distance of the nearest primary object and intensity gradients of cell membrane image [43]. Hereafter we refer these methods as CP-distance, CP-watershed, and CP-propagation, respectively. Results show that Dice-XMBD outperformed all other benchmarked methods with highest accuracy on pixel level (F1 score = 0.92, Jaccard index = 0.85) (Figure 2A). We also observed that CP-distance obtained the highest recall (Recall = 0.95) but lowest precision (Precision = 0.66), which means that it can identify almost every pixel correctly in the labeled mask but only 66% of predicted pixels were accurate.

In terms of cell-level performance, we first counted cells per image from predicted and labeled cell masks. Dice-XMBD can predict cells similar to ground truth (Pearson correlation = 0.998). Although the correlation between prediction and ground truth was relatively high among all segmentation methods, Mesmer tended to predict less cells while CellProfiler was more likely to over-split cells, as shown in Figure 2B and Figure 2C. Moreover, Figure 2C shows that Dice-XMBD had the best prediction performance (F1-score = 0.856) considering precision (Precision = 0.880, percent of cells that were correctly predicted) and recall (Recall = 0.867, percent of true cells that are predicted) than Mesmer (F1-score = 0.557) and CellProfiler (F1-score *≈* 0.56). We further checked the IOU distribution of all one-to-one cell pairs (predicted and true cells), Figure 2D demonstrates that most matched cell pairs predicted from Dice-XMBD were highly overlapping (mean = 0.815, median = 0.821), followed by Mesmer where most matched pairs are only half area of overlap (mean = 0.579, median = 0.595). An example of BRCA1 shown in Figure 2E demonstrates Dice-XMBD prediction was far superior to other benchmarked methods since it contained most cells with high matched values.

### 3.2 Dice-XMBD enables generic IMC image segmentation tasks

The key idea of this study was to generate an IMC-specific single cell segmentation model across different datasets with multiple proteins. We selected three independent IMC datasets generated from different labs to test the generalization ability of Dice-XMBD. Apart from the benchmarked methods mentioned above, we also included Ilastik model trained from BRCA1 annotations in our comparison. Figure 3A shows that Dice-XMBD outperformed all the methods, followed by Ilastik. Moreover, the performance of cells prediction from Dice-XMBD was the best and the most stabilized among three datasets, while Ilastik and Mesmer tended to under-predict cells. CellProfiler predicted less cells in BRCA2 and over-predicted cells in two T1D datasets, as shown in Figure 3B and Figure 3C. Furthermore, Dice-XMBD predictions contained most of the cells with IOU value higher than 0.8 (Figure 3D and Supplementary Figure S3).

**Figure 3:**
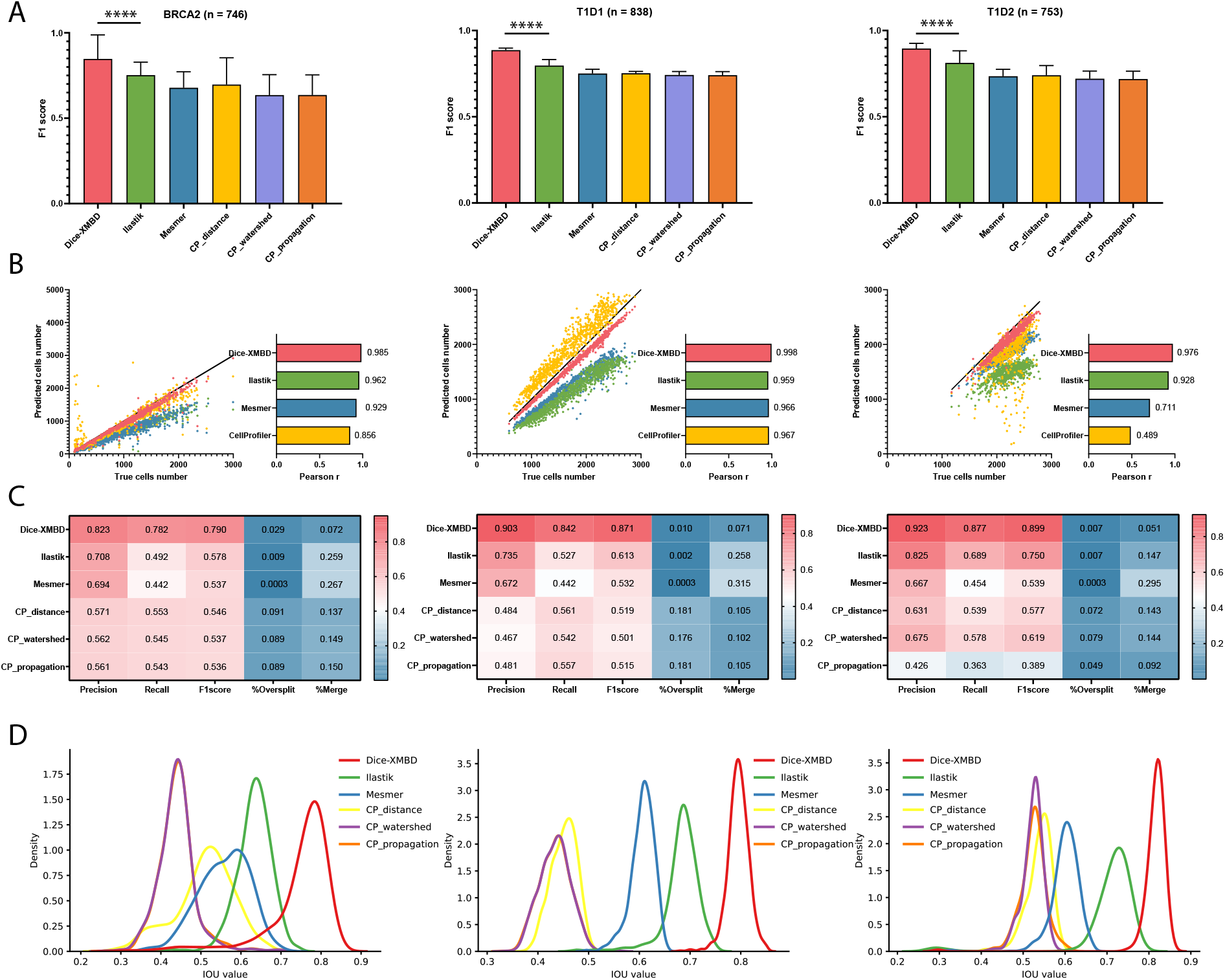
Dice-XMBD enables generic IMC image segmentation. Left: BCRA2, middle: T1D1, right: T1D2. **(A)** Pixel prediction performance comparisons of Dice-XMBD, Ilastik, Mesmer and CellProfiler (CP-distance, CP-watershed, CP-propagation). All data in bar plots are presented as mean values +/-SD. **(B)** Pearson correlations between the number of predicted cells and labelled cells per image. The number of cells predicted from three cell segmentation methods implemented in CellProfiler are the same (represented as CellProfiler). **(C)** Heatmaps of cells prediction performance of six benchmarked methods. The percent of oversplit and merge errors in predictions are denoted as %Oversplit and %Merge. **(D)** Density plots show the distribution of mean IOU values of matched cells per image.

### 3.3 Dice-XMBD enables accurate downstream biological analysis

Single cell segmentation is the first step of downstream analysis, such as protein quantification, cells clustering and annotation. Figure 4A shows the distribution of five proteins extracted from one image of BRCA1, proteins distribution from Dice-XMBD are more similar to them from ground truth than other methods. We calculated Pearson correlation of protein profiling between prediction and ground truth, which demonstrated that protein profiling correlation from Dice-XMBD was the highest among all methods by testing in the same dataset (BRCA1) and across datasets (BRCA2, T1D1, T1D2), as shown in Figures 4B and 4C. These results suggest that Dice-XMBD has good generalization ability to predict single cells for different IMC images with minimum impact to the downstream analysis due to the high correlation of its results with ground truth.

**Figure 4:**
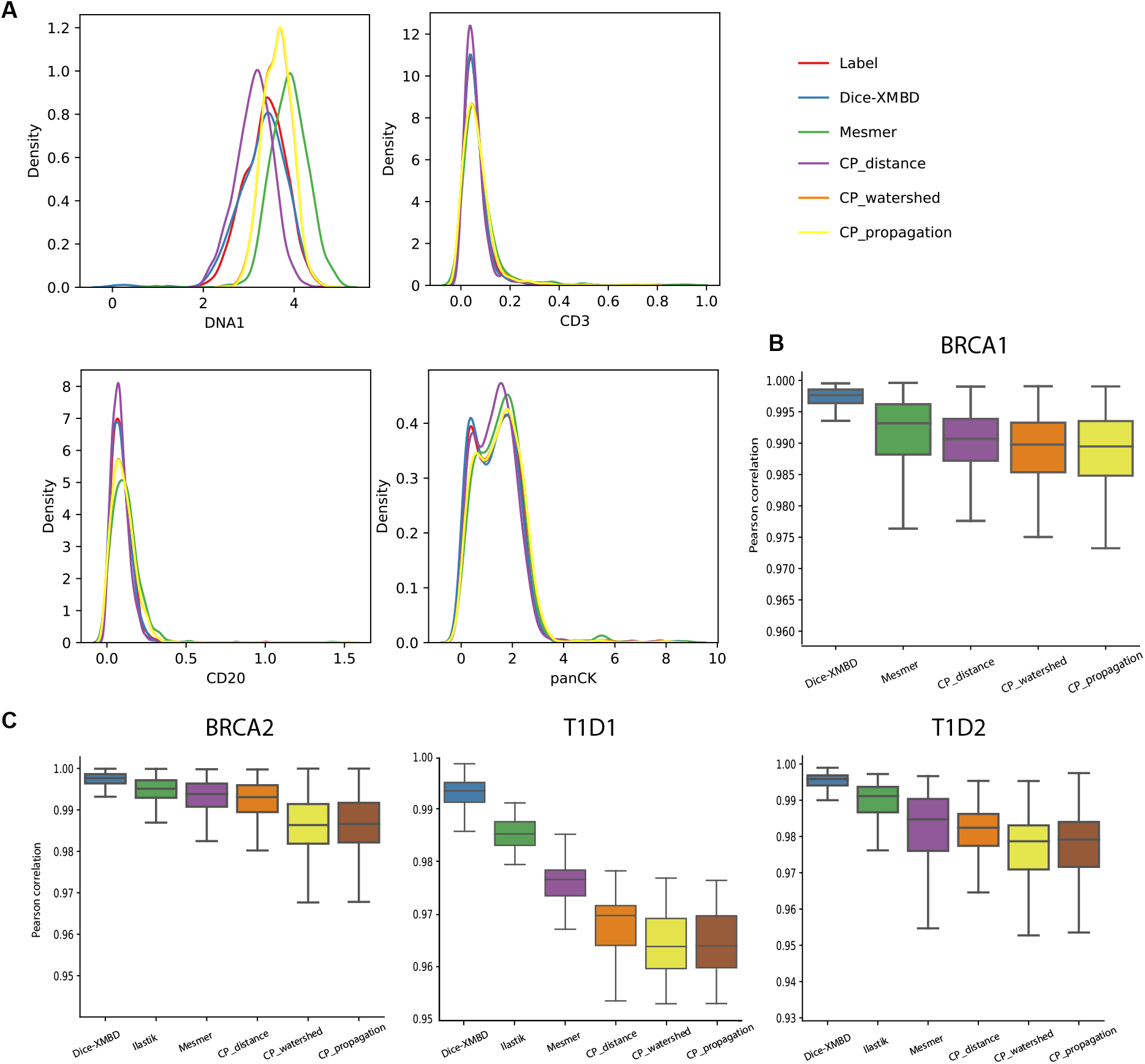
Dice-XMBD enables accurate downstream biological analysis. **(A)** Four protein distribution of one image of BRCA1. Sing cell protein profiling Pearson correlation between prediction and ground truth for BRCA1 dataset **(B)** and other three datasets **(C)**. Boxplots represent median with the center line, and whiskers extend to 1.5x interquartile range (IQR).

## 4. DISCUSSION

Highly multiplexed single cell imaging technologies such as IMC are becoming increasingly important tools for both basic biomedical and clinical research. These tools can unveil complex single-cell phenotypes and their spatial context at unprecedented details, which provide a solid base for further exploration in cancer, diabetes, and other complex diseases. Nevertheless, cell segmentation has become a major bottleneck in analyzing multiplexed images. Conventional approaches rely on intensities of protein markers to identify different cellular structures such as nuclei, cytoplasm, and membrane. Unfortunately, the intensity values of these markers are strongly cell type-specific and may vary from cells to cells. In addition, the staining also shows variability across images or datasets. As a result, the accuracy and robustness of the segmentation results are far from optimal. On the other hand, high-order visual features including spatial distribution of markers, textures, and gradients are relevant to visually identify subcellular structures by human. However, those features are not considered in these methods to improve the cell segmentation results.

The DNN-based image segmentation approaches provide an opportunity to leverage high-order visual features at cellular level for better segmentation results. Unfortunately, they require a significant amount of annotation data that are in general difficult to acquire. In addition, the highly variable channel configurations of multiplexed images impose another important obstacle to the usability of these methods as most of them lack the ability to adapt to different channel configurations after models are trained. In this study, we develop Dice-XMBD, a generic solution for IMC image segmentation based on U-Net. Dice-XMBD overcomes the limitation of training data scarcity and achieves human-level accuracy by distilling expert knowledge from Ilastik with manual input of human as a teacher model. Moreover, by consolidating multiple channels of different proteins into two cellular structure aware channels, Dice-XMBD provides an effective off-the-shelf solution for cell segmentation tasks across different studies without retraining that can lead to significant delay in analysis. Finally, to facilitate the analysis of large amount of IMC data currently being generated around the world, we made Dice-XMBD publicly available as an open-source software on GitHub (https://github.com/xmuyulab/Dice-XMBD).

## Author Contributions

WY, LW, RY and JH discussed the ideas and supervised the study. YQ and YJ implemented and conducted experiments in deep network cell segmentation. XX performed the model evaluation and biological analysis on segmentation results. XX, WY and RY wrote the manuscript. All authors discussed and commented on the manuscript.

## Data Availability Statement

All datasets used for this study can be found at GitHub (https://github.com/xmuyulab/Dice-XMBD). These datasets are downloaded from: BRCA1 (https://idr.openmicroscopy.org/search/?query=Name:idr0076-ali-metabric/experimentA), BRCA2 (https://zenodo.org/record/3518284#.YLnmlS8RquU), T1D1 (https://data.mendeley.com/datasets/cydmwsfztj/1), T1D2 (part1: https://data.mendeley.com/datasets/9b262xmtm9/1, part2: https://data.mendeley.com/datasets/xbxnfg2zfs/1), respectively.

