## Supplementary for "Dice-XMBD: Deep learning-based cell segmentation for imaging mass cytometry"

Xiao et al

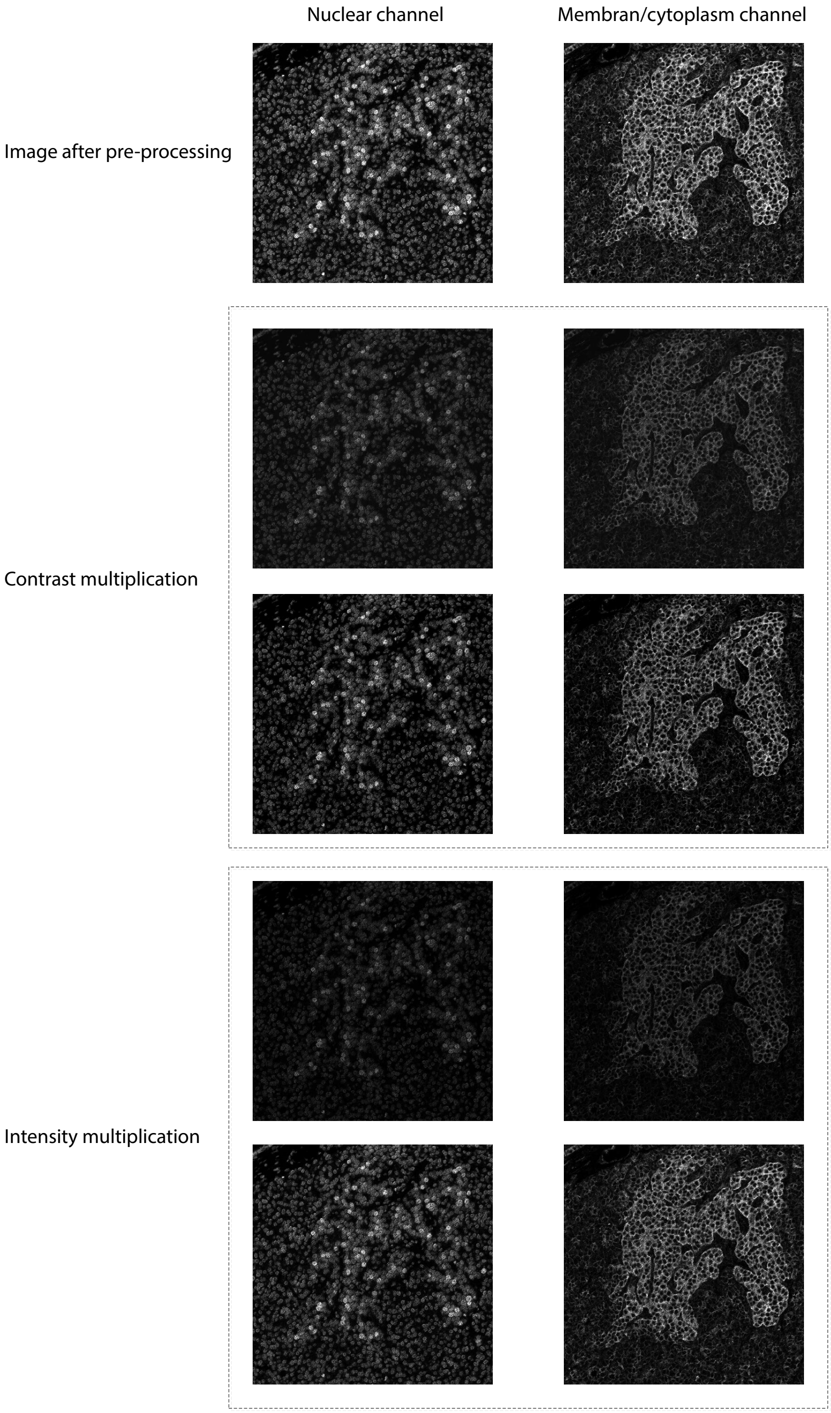


Figure 1: An example of data augmentation: 2-channel images after preprocessing, contrast and intensity multiplication.

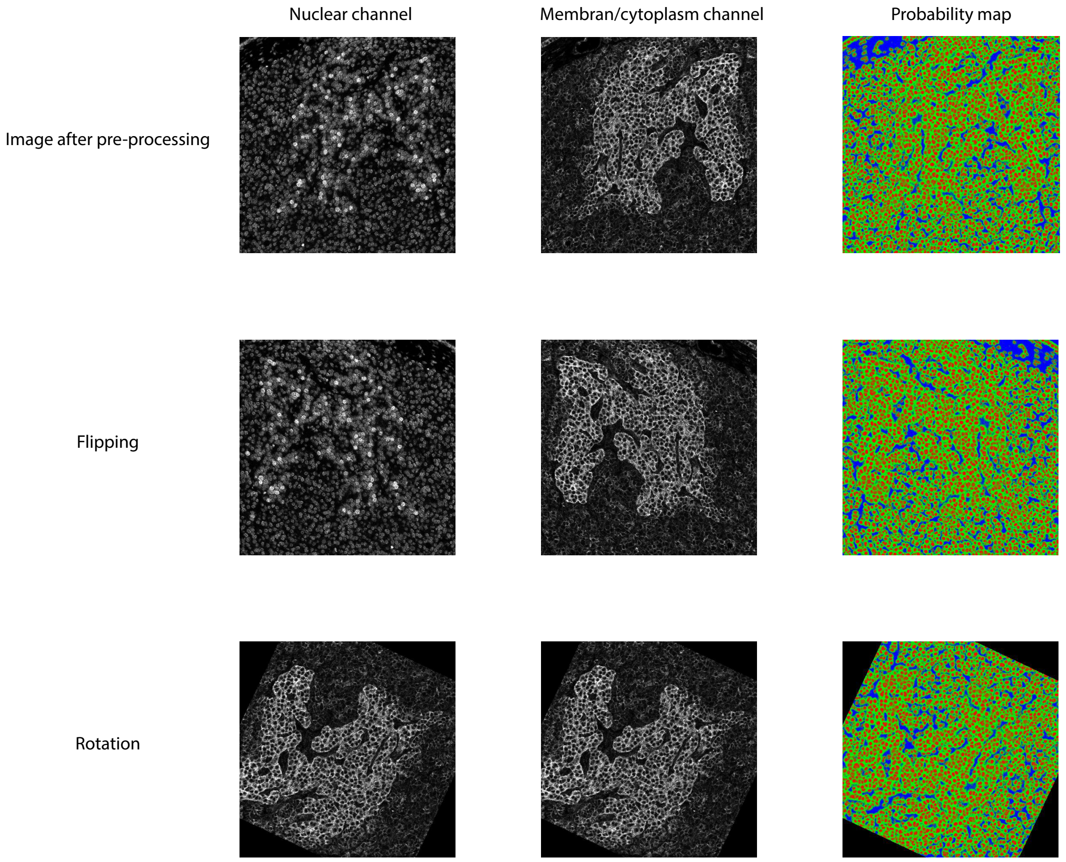


Figure 2: An example of data augmentation: 2-channel images and and probability maps after preprocessing, flipping, and rotation.

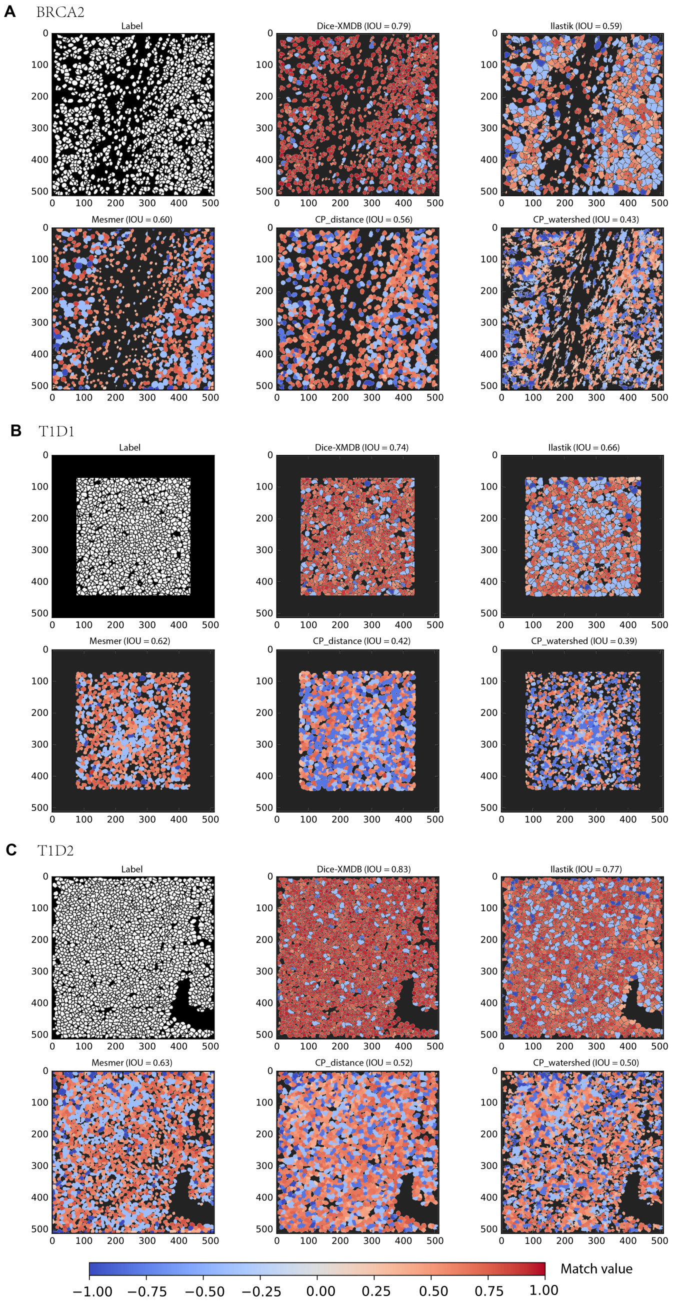


Figure 3: An example of labeled and predicted single cell mask from benchmarked methods for dataset BRCA2 **(A)**, T1D12 **(B)**, and T1D2 **(C)**. Match value represents IOU value for one-to-one cell pair found in label and prediction, -0.4 and -0.8 for merged cells (multiple true cells are assigned to one predicted cell) and split cells (multiple predicted cells are matched to a true cell), and -1 for cells are not in any of above situations. Numbers in brackets of each method indicate mean of IOU values of all matched cell pairs.
